# No one tool to rule them all: Prokaryotic gene prediction tool performance is highly dependent on the organism of study

**DOI:** 10.1101/2021.05.21.445150

**Authors:** Nicholas J. Dimonaco, Wayne Aubrey, Kim Kenobi, Amanda Clare, Christopher J. Creevey

**Affiliations:** Institute of Biological, Environmental and Rural Sciences, Aberystwyth University, Aberystwyth, SY23 3PD, Wales, UK; Department of Computer Science, Aberystwyth University, Aberystwyth, SY23 3DB, Wales, UK; Department of Mathematics, Aberystwyth University, Aberystwyth, SY23 3BZ, Wales, UK; School of Biological Sciences, Queen’s University Belfast, Belfast, BT7 1NN, Northern Ireland, UK

## Abstract

**Motivation:** The biases in Open Reading Frame (ORF) prediction tools, which have been based on historic genomic annotations from model organisms, impact our understanding of novel genomes and metagenomes. This hinders the discovery of new genomic information as it results in predictions being biased towards existing knowledge. To date users have lacked a systematic and replicable approach to identify the strengths and weaknesses of any ORF prediction tool and allow them to choose the right tool for their analysis.

**Results:** We present an evaluation framework (ORForise) based on a comprehensive set of 12 primary and 60 secondary metrics that facilitate the assessment of the performance of ORF prediction tools. This makes it possible to identify which performs better for specific use-cases. We use this to assess 15 *ab initio* and model-based tools representing those most widely used (historically and currently) to generate the knowledge in genomic databases. We find that the performance of any tool is dependent on the genome being analysed, and no individual tool ranked as the most accurate across all genomes or metrics analysed. Even the top-ranked tools produced conflicting gene collections which could not be resolved by aggregation. The ORForise evaluation framework provides users with a replicable, data-led approach to make informed tool choices for novel genome annotations and for refining historical annotations.

**Availability:** https://github.com/NickJD/ORForise

**Supplementary information:** Supplementary data are available at bioRxiv online.

## 1 Introduction

Whole genome sequencing, assembly and annotation is now widely conducted, due predominantly to the increase in affordability, automation and throughput of new technologies (Land *et al*., 2015). ORF prediction in prokaryote genomes has often been seen as an established routine, in part due to a number of assumptions and features such as the high density (protein coding genes contribute *∼*80-90% of prokaryote DNA) and the lack of introns (Lobb *et al*., 2020; Salzberg, 2019). However, this process involves the complex identification of a number of specific elements such as: promoter regions (Browning and Busby, 2004), the Shine–Dalgarno (Dalgarno and Shine, 1973) ribosomal binding site, and operons (Dandekar *et al*., 1998), which all contribute to identifying gene position and order. Additionally, the role of horizontal gene transfer (HGT) (Jain *et al*., 1999) and pangenomes further complicates an already difficult process and likely contributes to errors and a lack of data held in public databases (Devos and Valencia, 2001; Furnham *et al*., 2012). Finally, our ability to characterise the functions of regions of DNA (which has been generally reserved for model organisms and core genes (Russell *et al*., 2017)) is being outstripped by the rate of genomic and metagenomic sequence data generation from non-model organisms and non-core gene DNA sequences.

Before the turn of the century, it was understood that a great deal of work was still needed to address these issues. Studies had shown that many existing ORF prediction tools systematically fail to identify or accurately report gene families whose features lay outside a rigid set of rules, such as non-standard codon usage, those which overlap other genes or those below a specified length (Guigo, 1997; Burge and Karlinb, 1998). While there has been much work to address this, many gene families continue to be absent or underrepresented in public databases (Warren *et al*., 2010; Huvet and Stumpf, 2014), such as short/small-ORFs (short ORFs) (Storz *et al*., 2014; Duval and Cossart, 2017; Su *et al*., 2013). This means that ORF prediction methodologies that use information from existing sequences are in turn ill-equipped to identify genes belonging to these underrepresented/missing gene families. It is therefore of paramount importance that we understand the limits of current ORF predictors as our reliance on automated genome annotation continues to increase (Brenner, 1999). Measures to compare both novel and contemporary ORF prediction tools are not well established or universally employed and novel tool descriptions tend to focus on the algorithmic improvements rather than carrying out a systematic assessment of where the weaknesses in their approach lies. This prevents researchers from gaining meaningful insight into the specific features of genes which lead to them being missed or partially detected, resulting in a lost opportunity to improve our understanding of prokaryote genome content.

To address this, we extensively evaluate a collection of 15 widely used ORF prediction tools that form the basis of most of the annotations deposited in public databases and therefore have largely been used to build the genomic knowledge used by the scientific community. We provide a comparison platform developed to allow researchers to compare 12 primary and a further 60 secondary metrics to systematically compare the predictions from these tools and study the effect on the resulting genome annotations for their species of interest. This allows for in-depth and reproducible analyses of aspects of gene prediction that are often not investigated and allows researchers to understand the impact of tool choice on the resulting prokaryotic gene collection.

## Materials and Methods

### Gold standard genome annotations

Six bacterial model organisms and their canonical annotations were downloaded from Ensembl Bacteria (Howe *et al*., 2020)^1^. They were chosen for their phylogenetic diversity, scientific importance, range of genome size, GC content, assumed near complete and high quality genome assembly and annotation provided by Ensembl Bacteria. They are presented in detail in Table 1 and further information regarding these model organisms can be found in supplementary section 1.

**Table 1:** An overview of genome composition for the 6 model organisms selected to evaluate ORF prediction tools compiled from data held by Ensembl Bacteria. Data is presented for all genes and Protein Coding Genes (PCGs) in bold square brackets. Note the relatively broad differences in genome size, gene density (percentage covered with annotation) and GC content.

| Model Organism | Genome Size (Mbp) | Genes [PCGs] | Genome Density [PCGs] | GC Content |
| --- | --- | --- | --- | --- |
| *B. subtilis* | 4.04 | 4,133 [4,011] | 88.91% [87.60%] | 43.89% |
| *C. crescentus* | 4.02 | 3,875 [3,737] | 90.60% [90.23%] | 67.21% |
| *E. coli* | 4.56 | 4,257 [4,052] | 86.28% [84.35%] | 50.80% |
| *M. genitalium* | 0.58 | 559 [476] | 92.03% [90.62%] | 31.69% |
| *P. fluorescens* | 6.06 | 5,266 [5,178] | 84.75% [84.20%] | 60.13% |
| *S. aureus* | 2.76 | 2,556 [2,478] | 83.93% [82.76%] | 32.92% |

For each of the chosen model organisms, two data files were downloaded from Ensembl Bacteria; the complete DNA sequence (*\* _dna*.*toplevel*.*fa*) and the GFF (General Feature Format) file (*\**.*gff3*) containing the position of each gene. The protein coding genes (PCGs) presented in the model organism annotations from Ensembl were taken as the gold standard (Ensembl Gold Standard - EGS) for this study. Bacterial genomes exhibit high levels of gene density, often with little extraneous DNA, which is “commonly perceived as evidence of adaptive genome streamlining” (Sela *et al*., 2016). Unannotated DNA represents between *∼*10%-20% of the six genomes selected and while an additional 0.38% - 2.22% is attributed to non-coding annotations, there is still a measurable portion of each genome without any annotation. This study focuses specifically on the identification of PCGs, which constitute the significant majority of annotated genomic regions in the bacteria studied (82.76% - 90.62%, see Fig 1).

**Figure 1:**
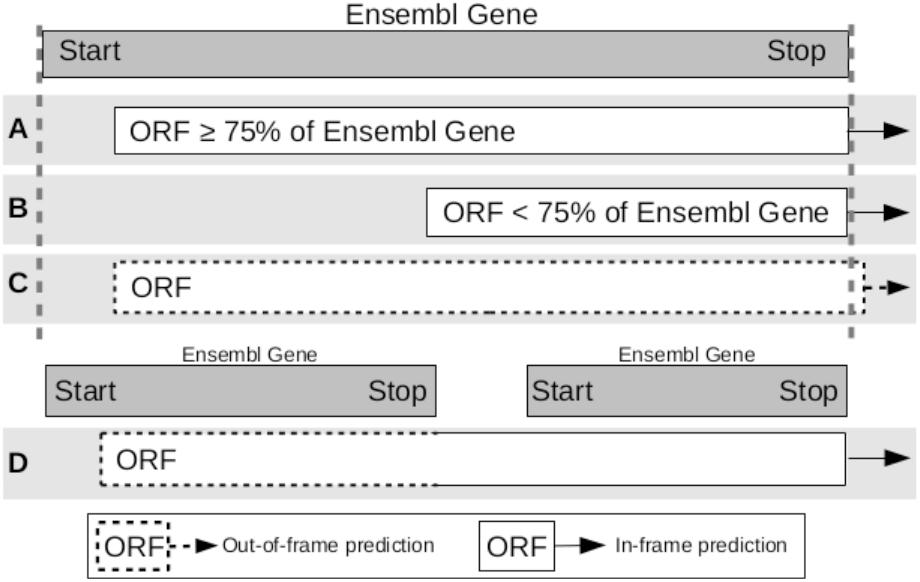
Illustration of how predicted ORFs are classified as having detected or not detected the EGS genes. Predicted ORFs are compared to the genes held in Ensembl. A - The predicted ORF covers at least 75% and is in-frame with Ensembl gene and therefore it is recorded as detected. B - The predicted ORF covers less than 75% of the Ensembl gene and therefore is recorded as not detected. C - The predicted ORF covers part of an Ensembl gene but is out of frame (dotted outline) and therefore is recorded as missed. D - The use of alternative stop codons causes the predicted ORF to be truncated or divided into two ORFs that span the Ensembl genes and therefore is recorded as missed.

The PCGs from each of the 6 model organisms exhibit a range of differences which are known to impact the ability of prediction tools to identify them. These include, but are not limited to, GC content, codon usage and gene length. The GC content varies from 31.69% - 67.21% for these genomes, and even within a single genome, the PCG GC content varies widely (see Supplementary Fig 1 for distributions). Furthermore, the canonical ATG start codon is used between 68.58% - 90.67% of the genes for the six genomes (see Supplementary Table 1 for more detail).

Additionally, *M. genitalium* uses the codon translation table 4, meaning one of the three universal stop codons (TGA/UGA) is instead used to code for tryptophan (Dybvig and Voelker, 1996), whereas the other 5 model organisms use the universal translation table 11 (see Supplementary Table 2 for more detail). While a similar median PCG length is shared across the six genomes, *B. subtilis* and *P. fluorescens* have a number of long genes (*>* 8,000 nt, see Supplementary Figure 2) and *S. aureus* contains the 31,421 nt “giant protein Ebh” (Cheng *et al*., 2014) which is more than twice the length of the next largest PCG in this study.

The Sequence Ontology (Eilbeck *et al*., 2005) describes an ORF as “The in-frame interval between the stop codons of a reading frame which when read as sequential triplets, has the potential of encoding a sequential string of amino acids”. However, it is the norm for ORFs to be reported as regions of DNA encompassed by a start and stop codon as a start codon is expected to indicate the start of DNA transcription (Brent, 2005). We acknowledge the difference in ontological definition and during this study, we refer to an ORF as the region of DNA between an in-frame start and stop codon which is predicted to encode for an amino acid (protein) sequence.

### Prediction tools

This study specifically investigates ORF predictors, tools which apply complex filtering after the identification of ORFs across a region of DNA. This is different to ORF finders, which return unfiltered ORFs (Stothard, 2000) that meet a set of pre-defined rules such as length and in-frame start and stop codons. This filtering is unique to each tool and dependent on properties such as codon usage, GC content, ORF length, overlap and similarity to known genes, and other more sophisticated parameters modelled on analysis of previously studied genes and genomes. Without such filtering methods, ORF finders would typically report many false positives such as nested or heavily overlapping ORFs. GeneMark (Borodovsky and McIninch, 1993) reports multiple variations of the same ORF with confidence scores and we chose the longest for each ORF after assessing the results.

We selected 15 different ORF prediction tools, some of which required a model (a rigid set of parameters pre-tuned to a particular organism), and the others which predicted *ab initio* from sequence. The tools which required a model were: GeneMark.hmm with *E. coli* and *S. aureus* models (Lukashin and Borodovsky, 1998); FGENESB with *E. coli* and *S. aureus* models (Salamov and Solovyevand, 2011); Augustus with *E. coli, S. aureus* and *H. sapiens* models (Keller *et al*., 2011); EasyGene with *E. coli* and *S. aureus* models (Nielsen and Krogh, 2005); GeneMark with *E. coli* and *S. aureus* models (Borodovsky and McIninch, 1993). Those which did not require a model were: GeneMarkS (Besemer *et al*., 2001); Prodigal (Hyatt *et al*., 2010); MetaGeneAnnotator (Noguchi *et al*., 2008); GeneMarkS-2 (Lomsadze *et al*., 2018); MetaGeneMark (Zhu *et al*., 2010); GeneMarkHA (Besemer and Borodovsky, 1999); FragGeneScan (Rho *et al*., 2010); GLIMMER-3 (Delcher *et al*., 2007); MetaGene (Noguchi *et al*., 2006); TransDecoder (Haas *et al*., 2013). Included in this list is a number of tools which were designed for fragmentary and metagenomic studies: MetaGeneMark, MetaGene, MetaGeneAnnotator and FragGeneScan. In addition, TransDecoder was developed to predict coding regions within transcript sequences, often in eukaryotes. The two groups are referred to as ‘model-based’ and ‘*ab initio*’ henceforth. To emulate the annotation process of a novel or less studied genome or metagenome, each tool was run using its default parameters. More information regarding each group and tool, and the parameters used to run them, can be found in Supplementary Section 3 Prediction Tools.

Whole genome annotation ‘pipelines’ such as PROKKA (Seemann, 2014) and NCBI’s PGAP (Tatusova *et al*., 2016) were not included, but the ORF prediction components embedded in these pipelines such as Prodigal and GeneMarkS-2 were included in the study. Multiple separate tools from the GeneMark family (Besemer and Borodovsky, 2005) were included (some superseded) due to their extensive use and impact on genomic knowledge over the last three decades.

### Comparison method

A systematic software platform ORForise (ORF Authorise) was built to perform a fair, comparative, and informative analysis of the different tools examining different aspects of their predictions. Version 1.0 of the platform, written in Python3 (Van Rossum and Drake, 2009), was used and is freely available at https://github.com/NickJD/ORForise. It has been designed to process the standardised GFF3 format as well as the individual output formats produced by each tool listed in this study.

In this platform we endeavoured to choose a wide range of metrics that clearly and representatively capture the many intricacies of the predictions. A number of metrics used in previous studies, such as the number of ORFs predicted, accurate identification of start positions or the number of genes correctly detected, can give some indication of the ‘accuracy’ of each tool. However, it was found during our analysis that there were many complexities in the prediction results which would not be represented by these high-level metrics. For example, predicted ORF regions may overlap with one or more known EGS genes but be inaccurately extended or truncated on either the 5’ or 3’ end. It is also common for smaller EGS genes to be mistakenly encompassed by larger predicted ORFs and while the nucleotide regions of these genes are technically within the predicted regions, even if in-frame, they do not represent the true protein coding sequence. Therefore, clear and specific measures of accuracy that describe the detection of the entire locus of a gene are needed. Figure 1 illustrates how we determine correct EGS gene detection, but also explains its nuances and complexities. An example of this is the definition of short ORFs, which in bacteria are often described as having lengths of 100-300 nt (Storz *et al*., 2014; Duval and Cossart, 2017; Su *et al*., 2013). However, due to hard-coded cutoffs in many of the tools, we chose the ‘upper-bound’ of 300 nt or 100 codons to define short ORFs.

We iteratively developed 72 metrics to help provide the most accurate and informative representation of a tool’s prediction quality. Additionally, as part of the ORForise platform, we provide a number of Python3 post analysis scripts developed to aid in the interrogation between the EGS gene annotations and the ORFs predicted by each of the tools studied. These scripts were used to extract characteristics that are useful in the investigation of why specific EGS-genes are detected or missed.

### Aggregated tool predictions

An extension to the ORForise comparison platform was built to investigate whether an aggregation of predictions from a number of top-performing tools would perform better than individual tools. The ORF predictions from the selected tools are combined into a single data structure with duplicate ORFs filtered out, but alternative predictions for the same locus retained and ordered according to start position. The same comparison algorithm could then be employed on the set of unique ORF predictions identified by this union of the outputs of the selected tools (Prodigal, GeneMark-S-2, MetaGeneAnnotator, MetaGeneMark and GeneMark-S - chosen due to their individual performance) and as with the singular comparison, for every EGS-gene, the ORF which deviated the least from the correct locus was selected as the closest match.

### Discovering additional ORFs

To enable the aggregation of different Gold Standard and predicted ORFs, we provide GFF_Intersector to create a single GFF representing the intersection of two existing GFFs. This also provides an option to allow the retention of genes that have a user defined difference (default minimum 75% coverage and in-frame).

To enable the addition of predicted ORFs to an existing GFF, we also provide the GFF_Adder tool, which produces a new GFF containing the original genes plus the new ORFs, filtered to remove any ORFs that overlap existing genes by more than 50 nt (user definable).

## Results

### Metrics for comparison of tools

72 different metrics were chosen for this exhaustive evaluation in order to give the broadest possible scope to compare and contrast the performance of the tools. The full definitions for each of these metrics can be found in Supplementary Section 5 and are intended to be used as a resource for the community when deciding which tool to apply to both novel and contemporary genome annotation work. The following are 12 of the most informative metrics, selected for their ability to represent both a broad range and depth of different attributes which have been used to distinguish the prediction tools.

- **M1** Percentage of Genes Detected
- **M2** Percentage of ORFs that Detected a Gene
- **M3** Percentage difference of number of Predicted ORFs
- **M4** Percentage difference of Median ORF Length
- **M5** Percentage of Perfect Matches
- **M6** Median Start Difference of Matched ORFs
- **M7** Median Stop Difference of Matched ORFs
- **M8** Percentage difference of Matched Overlapping ORFs
- **M9** Percentage difference of Matched Short ORFs
- **M10** Precision
- **M11** Recall
- **M12** False Discovery Rate

For M3, M4, M8 and M9, *Percentage Difference* was used to identify differences between predicted and gold standard metrics: 100 *\*(ORF metric - Ensembl Gene Metric) / Ensembl Gene Metric*. The best score for a metric using the *Percentage Difference* calculation is 0, as 0 represents no deviation from the EGS annotations. The ‘Matched ORFs’ identifier used for M6, M7, M8 and M9 represent the ORFs which have correctly detected an EGS gene. M6 and M7 are calculated by taking the median codon position differences recorded for mispredicted start or stop codons. Metrics such as the Percentage of Perfect Matches (M5) can give a clearer overview of a tool’s ‘accuracy’ or performance, as it is common for a tool to misidentify either the exact start or stop locus of a detected gene, while metrics such as Median Start Difference of Matched ORFs (M6) can help establish the level of inaccuracy.

The tools were ordered by totalling the rankings for each of these 12 metrics, across the 6 model organisms. Supplementary Results 1 contains the results used for the ranking.This ranking, based on a wide range of different performance measures, allows for a comparative overview of contemporary and future tools, and is presented in Figure 2. This figure also shows the Percentage of Genes Detected (M1) with an overlay of the Percentage of Perfect Matches (M5), demonstrating the inconsistency between the two metrics for each tool. Metrics such as Percentage of Genes Detected (M1) and Percentage of ORFs that Detected a Gene (M2) are informative and can be representative of a tool’s prediction quality, however, they do not convey the complete picture when presented in isolation. This is of particular importance for those working with metagenomic or other fragmentary assemblies, as the likelihood of incomplete fragments and chimeric sequences is higher and can lead to varying mispredictions. Although the overall prediction quality of genes was high across most of the tools and genomes in this study, the additional metrics produced can be used to identify strengths and weaknesses inherent to them. For example, GeneMark.hmm (*S. aureus* model and genome), MetaGeneMark and MetaGeneAnnotator, GeneMarkS were all ranked highest for Percentage of Genes Detected (M1) for at least one model organism, while Prodigal and GeneMarkS were ranked highest twice. However, when inspecting the 12 chosen metrics (Figure 3), it was clear that there were complex differences between the prediction results of not only the highest scoring tools, but also the lower ranked tools which were often ranked high for some metrics.

**Figure 2:**
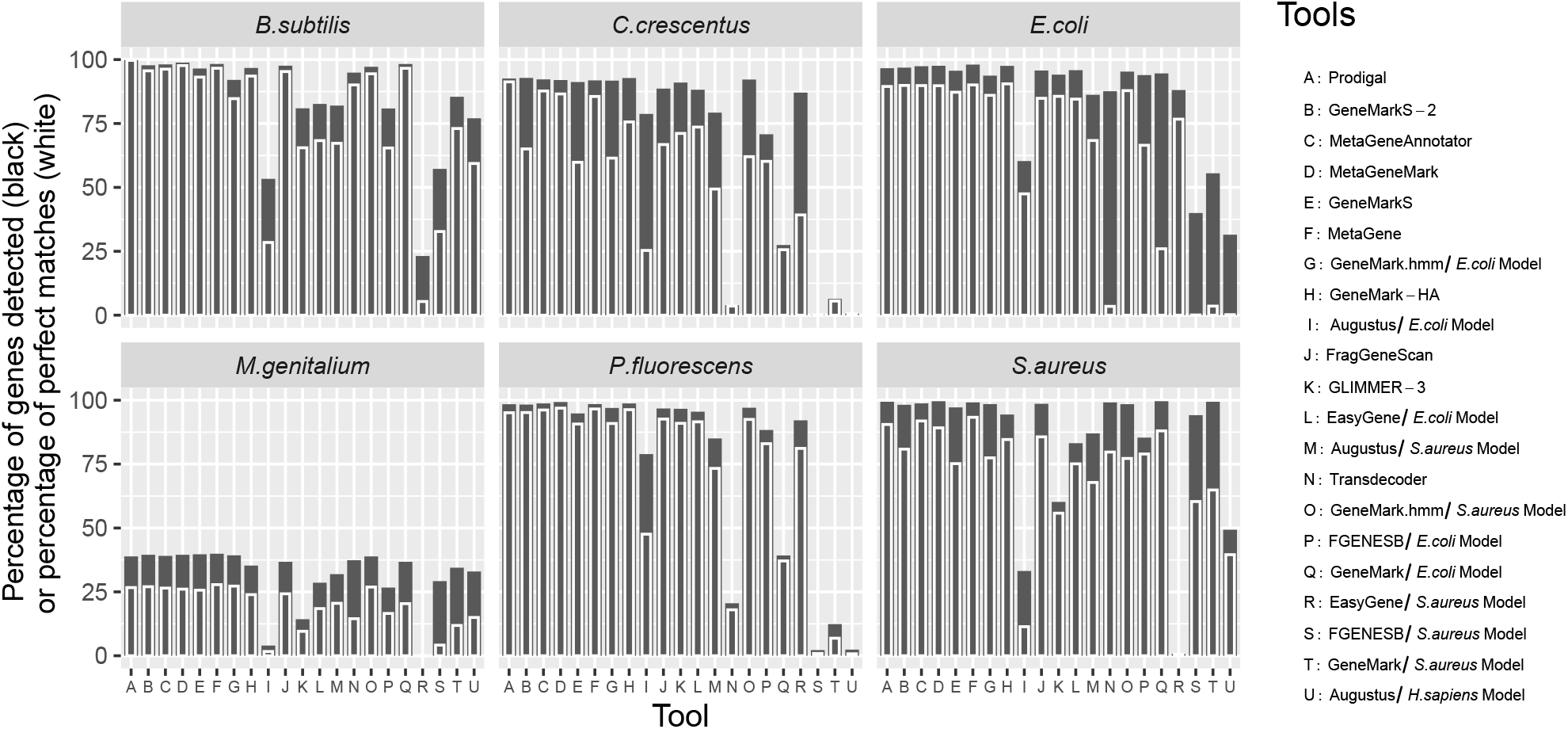
The result of all 15 gene prediction tools (21 with chosen models) on the 6 model organism genomes, ordered by the summed ranks across the 12 metrics. The *Y* axis represents the Percentage of Genes Detected (M1) by each tool in black and the Percentage of Perfect Matched (M5) in white. M5 which represents the ability for a tool to detect the correct start codon, has more variance between the tools than M1. Each column on the *X* axis represents a different tool (some model based tools were run multiple times). There is considerable variation in how well each tool performs across the different genomes, while all tools perform relatively poorly on the *M. genitalium* genome.

**Figure 3:**
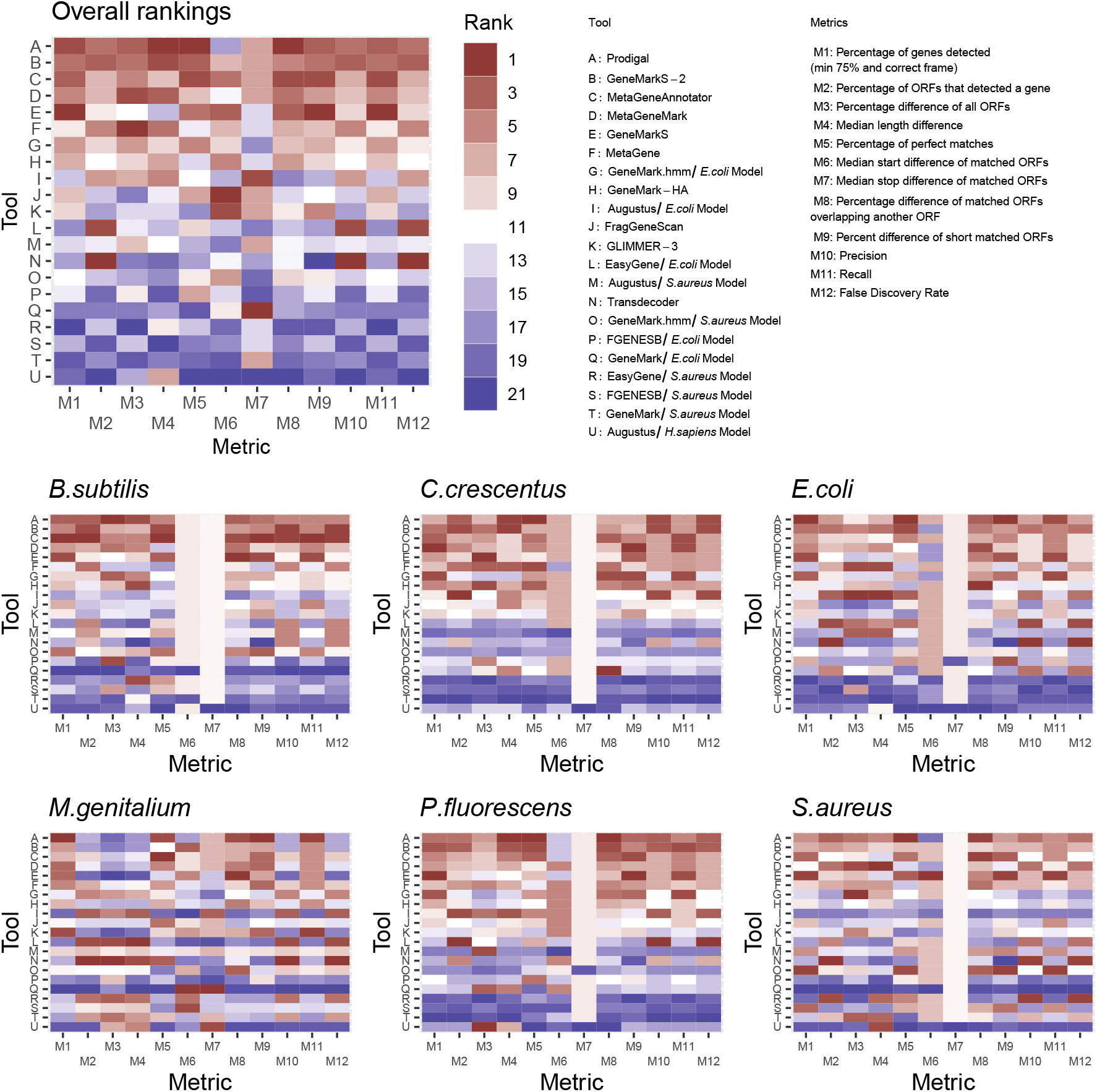
Heatmaps showing rankings of the tools by the 12 chosen metrics, overall and for each organism in turn. The tools are shown ordered by the summed ranks across the 12 metrics. While red is ‘better’ and blue is ‘worse’, it is clear that across the 6 model organisms, no tool stands out for these 12 metrics chosen as most representative. For example, for *C. crescentus*, GeneMark with *E. coli* model ranked 12th overall but reported the most accurate number of overlapping Genes. For *P*.*fluorescens*, Prodigal was the overall highest ranked tool even though GeneMarkS detected the highest number of Ensembl genes. *M. genitalium* on the other hand, which uses an alternative stop codon, has some very interesting results showing the difficulty to identify it’s genes by all tools. The pale coloured bands represent tools ranking the same for a particular metric.

While no tool or group of tools were consistently ranked highest or equally across the 12 metrics or model organisms, MetaGeneAnnotator ranked best for *B. subtilis* and *M. genitalium*, GeneMarkS-2 ranked best for *C. crescentus* and Prodigal ranked best for *E. coli, P. fluorescens* and *S. aureus*.

The combination of multiple metrics can be used to determine which tool should be used between two candidate tools with the same or similar Percentage of Genes Detected (M1), such as GeneMarkS and MetaGeneMark, which when applied to *M. genitalium* both obtained an M1 score of 39.50%. Meta-GeneMark reported a higher Percentage of Perfect Matches (M5) (65.96% compared to 61.17%) than GeneMarkS, as can be seen in Figure 2. In addition, GeneMarkS is ranked first for Percentage of Genes Detected (M1) when applied to *P. fluorescens* with 99.29%, compared to Prodigal which is ranked 4th with 98.49%. However, Prodigal has the highest Percentage of Perfect Matches (M5), 92.86% vs 87.03% for GeneMarkS, which means that more of the genes identified by Prodigal were exact matches. In this instance, choosing either Prodigal or GeneMarkS as the overall highest performing tool is not arbitrary.

### Model-based vs *ab initio* tools

It was evident that the performance of model-based tools was less consistent across the 6 model organisms than the *ab initio* tools. While they could perform as well as or better than a number of *ab initio* tools when the model selected was the same as the genome annotated, when it was not, they often produced predictions of extremely low quality. For example, GeneMark with the *E. coli* model only predicted 71 ORFs for *S. aureus’s* 2,478 genes, of which only 18 ORFs detected an EGS-gene. However, while it could be expected that mixing different models and genomes could cause poor quality predictions from model-based tools, there were instances in which both model and genome were the same and the prediction performance was also poor. In particular, in the case of EasyGene using the *S. aureus* model, only 49.31% of *S. aureus* EGS-genes were detected, a contrast from the *∼*99% detected by the majority of *ab initio* tools.

Intriguingly, Augustus (a model-based tool) when employed with the *E. coli* model, was able to detect 96.64% of *P. fluorescens* genes. While this shows that model-based tools can perform well even when their model and target genomes are different, when Augustus was applied to the same genome using the *S. aureus* model, it was only able to detect 20.53%, but unexpectedly detected 78.91% when using an *H. sapiens* model. This is indicative of the inconsistency of model-based prediction tools and the species-models they employ. In contrast, through the ranking approach we employed, the model-based tool GeneMark.hmm with the *E. coli* model ranked higher (7/21) than a number of *ab initio* tools in both the overall ranking and for individual metrics. Furthermore, GeneMark.hmm with the *S. aureus* model was joint top in detecting the highest number of *S. aureus* EGS-genes with GeneMarkS. Additionally, for each of the model-based tools, the *E. coli* model performed better across the 6 model organisms than the *S. aureus* model.

### GC content

No significant variation was observed between the EGS-gene median GC content and that of the predicted ORFs from each tool, even for those with poor predictions (see Supplementary Results File 2). This is likely due to the median GC content of the genomes being the driving factor for GC in any region of DNA, as the majority of the genomes are protein coding. However, when inspecting the results from Prodigal, some level of variability was observed in the different sets of Ensembl genes according to whether they were detected, partially matched, or missed, as can be seen in Supplementary Table 3. *E. coli* and *P. fluorescens* genes which were missed by Prodigal are nearly 10% lower in GC content than both detected and partial genes.

### Overlapping ORFs

The overall number of ORFs predicted to have an overlap with another ORF varied across each of the tools and model organisms, with cases of both positive and negative percentage differences when compared to the EGS annotations (see Supplementary File 2 ‘Full Prediction Metrics’). Proportionally, the number of overlapping ORFs reported by *ab initio* tools are closer to the number of EGS overlapping genes than those reported by the model-based group.

Most model-based tools underpredict the proportion of overlapping ORFs with the exception of GeneMark *E. coli* for *P. fluorescens*, which predicted 2,073 overlapping ORFs compared to the 1,251 reported by Ensembl (see Supplementary Table 4 and Results files 1 & 2).

Correct detection of EGS overlapping genes is also a problem. By totalling and averaging the Percentage Difference of Matched Overlapping ORFs (M8), we were able to observe a clear difference between the two tool groups with respect to their ability to detect correct overlapping EGS genes (see Supplementary Tables 4 and 5). The inability of the tools to account for the unusual nature of the *M. genitalium* genome was shown again with an average M8 across all tools of −88.21%, compared to the average of −27.77% for the other 5 genomes.

Furthermore, when making predictions for *E. coli*, while model-based tools such as Augustus and EasyGene with the *E. coli* model can closely predict the proportion of overall overlapping ORFs (Percentage Difference of −1.42% and −2.30% respectively), due to the poorer performance of these tools for correctly detecting EGS genes, their M8 scores for matched overlapping ORFs were substantially lower than the average score of the *ab initio* tools (grouped average of −52.89% as opposed to −23.62% - see Supplementary Table 5). Prodigal exemplifies this difference between the two tool groups. It was able to predict all overlapping EGS from *P. fluorescens* and *S. aureus*, whereas even when paired with the same model and genome, model-based tools continued to perform poorly.

### Short ORFs

Figure 4 summarises the gene lengths of detected, partially detected and missed genes when predicted by Prodigal. It shows that the EGS genes which were missed by Prodigal for each genome were substantially shorter in length than the genes which were detected (apart from *M. genitalium*). However, for the other 5 model organisms, whose combined median length of missed genes is 317, less than half the combined median length of 837.5 of those detected (Fig 4), it is alternative start codon selection which influences whether a predicted EGS is Shortened or elongated.

**Figure 4:**
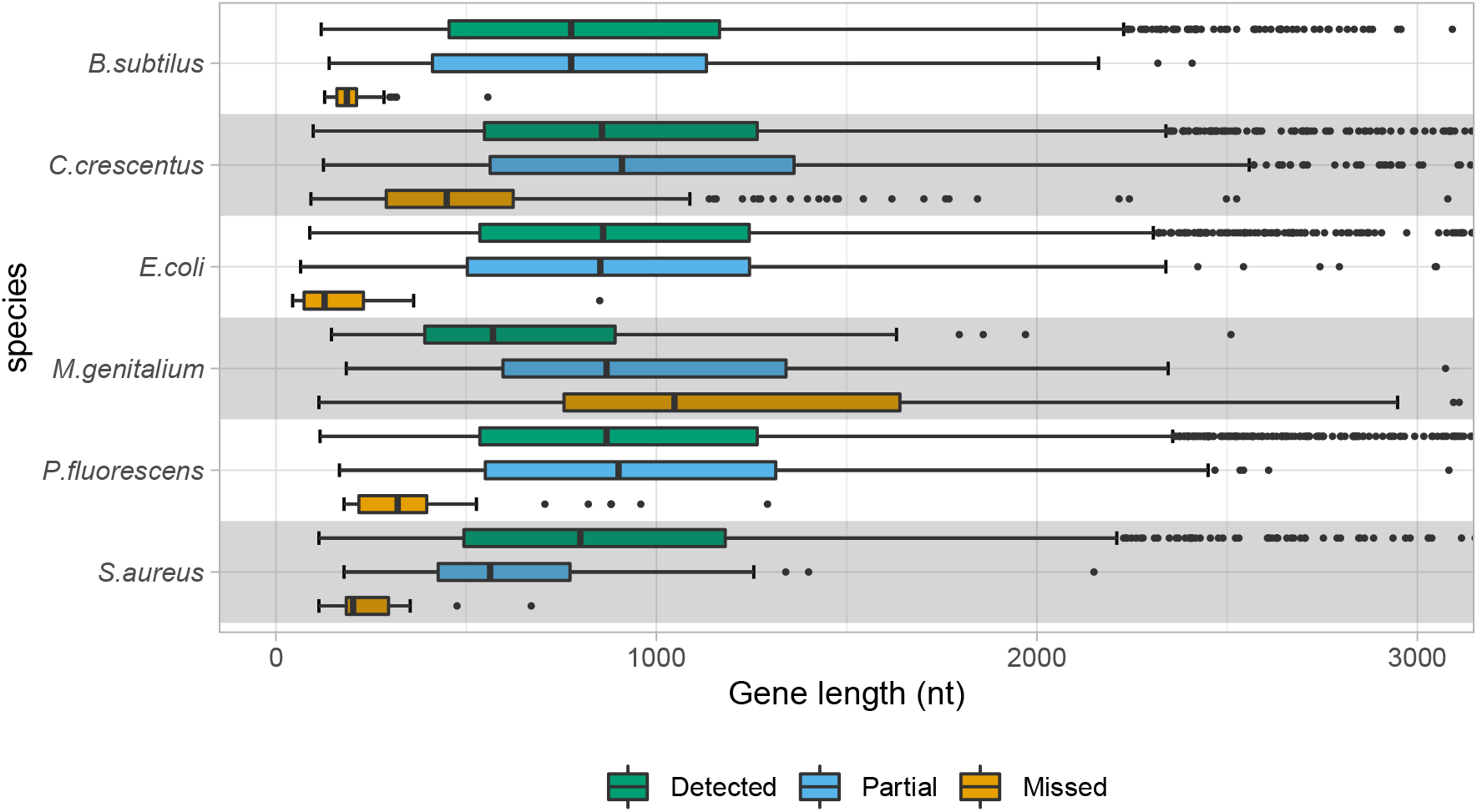
Lengths of Ensembl annotated genes, those which were partially detected by Prodigal and those which were missed, for each model organism. The x-axis is truncated at 3,000 nt. With the exception of *M. genitalium*, the distributions of lengths of the missed genes are generally to the left of the distributions of the detected genes. Thus Short genes are commonly overlooked by Prodigal and other tools.

The proportion of Short genes in the six Ensembl genomes below 300 nt ranged from 4.8% to 13.6% for each of the 6 model organisms. All tools predicted many short ORFs for *M. genitalium* because they were incorrectly truncated due to its alternative stop codon usage. On average, *Ab initio* tools were shown to be more likely to correctly detect EGS Short genes across the other 5 model organisms (see Supplementary Tables 4 and 5). Interestingly, unlike overlapping ORFs, short ORFs were more often overpredicted but few were actually accurate when compared to the EGS. However, *ab initio* tools were much better suited to reporting the correct proportion of short ORFs for all 6 genomes, often reporting the same proportion (see Supplementary Table 5). While *M. genitalium* does exhibit the highest divergence in proportional reporting of short ORFs, *ab initio* tools were still less divergent.

### Partial matches

The number of missed genes was low across the tools studied, with the exception of *M. genitalium* and some outliers from the model-based tools such as GeneMark, Augustus and EasyGene. However, we also found many genes that were incorrectly reported on the 5’ or 3’ end. These misannotations, which we have called ‘partial matches’ if in the correct frame and accounting for *>*= 75% of a gene, constitute either an elongation or truncation of the protein product of the gene and therefore potentially have an unknown impact on the resultant sequence. A large number of genes were incorrectly reported on the 3’ end for *M. genitalium* by each tool. These 3’ prime truncated ORFs are explained by the alternative use of TGA as a stop codon (normally used to encode tryptophan). The stop codons predicted for *M. genitalium* by all the prediction tools were the same ‘TGA,TAG,TAA’ as for the EGS-genes of the other five model organisms.

Unlike 3’ prime misprediction, a large number of genes from all six genomes were predicted with alternative start codons (see Supplementary Results File 2). This was true for all tools and especially a problem for *C. crescentus* with a relatively low 68.58% ATG start codon usage for all EGS-genes. The genes for which Prodigal was unable to obtain a ‘Perfect Match’ (M5), was just 37.40%. Prodigal used a much higher level of ATG (80.87%) for this set of partially detected genes. This misidentification of start codon usage was a consistent problem among all the tools and genomes studied. However, for *E. coli*, the level of misidentification was lower. Table 2 shows, as an example, the number of times the correct or incorrect start codons were selected by Prodigal, across all six model organisms, including the number of incorrectly chosen instances of the start codon (e.g. a different ATG further upstream of the real ATG).

**Table 2:** Start codon substitution table for genes which were misreported on the 5’ prime end by Prodigal, combined for the six model organisms. Column headers represent Ensembl annotated start codons and row headers represent the incorrectly predicted start codons, having chosen an alternative further upstream or downstream of the true start codon. The last row, ‘Correct codon’, shows the numbers of Perfect Match ORFs by Prodigal with the specified start codons. Further start codons with low usage were combined into the category labelled ‘other’.

|  | Correct codon |  |  |  |  |
| --- | --- | --- | --- | --- | --- |
|  | ATG | GTG | TTG | CTG | Other |
| Incorrect ATG | 817 | 371 | 357 | 19 | 24 |
| Incorrect GTG | 106 | 76 | 49 | 4 | 0 |
| Incorrect TTG | 81 | 47 | 37 | 3 | 3 |
| Incorrect CTG | 0 | 0 | 0 | 0 | 1 |
| Incorrect other | 0 | 0 | 0 | 0 | 0 |
| Correct codon | 14933 | 1321 | 847 | 0 | 0 |

### Aggregated tool predictions

Combined prediction approaches have previously been utilised to harness the prediction power from multiple tools to increase the number of genes detected (Yok and Rosen, 2011; Tatusova *et al*., 2016). For each of the model organisms, taking the union of the top 5 tool predictions did provide a small increase in the number of Genes Detected (M1) (and a reduction of partial matches) compared to that of the ‘best tool’ (tool with highest percentage of Genes Detected (M1)) for any particular organism. However, even with this extreme case of using the union of all tool predictions, the increase in M1 was negligible (average increase of 0.47%) and came at the expense of predicting a large number of additional incorrect ORFs, as can be seen in Table 3. Even in the case of *M. genitalium*, the M1 was not improved more than 0.21% with the union prediction.

**Table 3:**
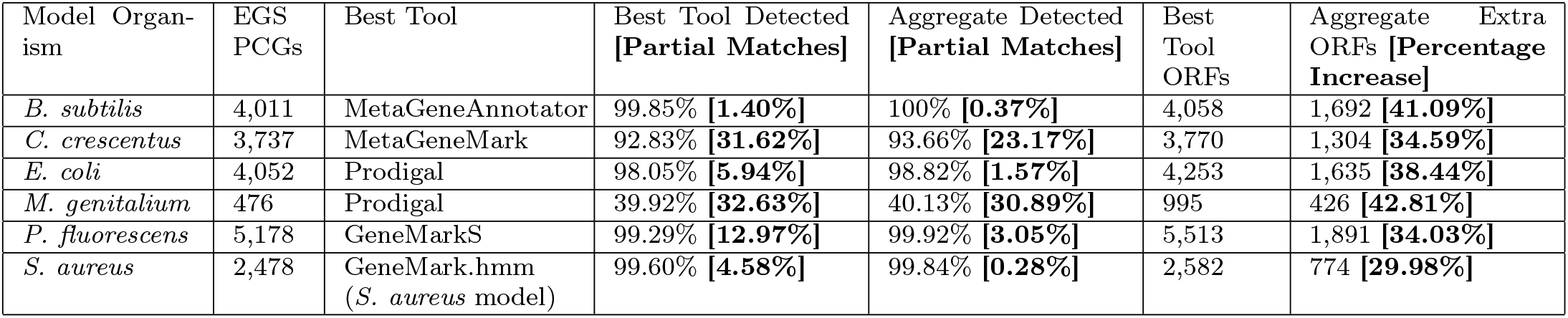
Aggregated tool predictions provide a small increase in Percentage of Genes Detected (M1) but over-predict a large number of additional ORFs. Here we compare the ‘best tool’ (tool with highest M1 score) predictions versus ‘aggregated tools’ (combination of predictions from top 5 ranked tools; Prodigal, GeneMark-S-2, MetaGeneAnnotator, MetaGeneMark and GeneMark-S) for the percentage of detected genes, partial matches and additional over-predictions made by the aggregated tools which did not detect an Ensembl Gold Standard (EGS) gene. GeneMark.hmm results are reported for *S. aureus* as even though it performed joint best with GeneMarkS (M1), it reported a higher proportion of Perfect Matches (M5).

### Improving historic annotations

Using the GFF_Adder tool, we investigated the the potential of Prodigal to add additional ORFs to the EGS annotations. Table 4 shows that there are more than 60 additional predicted ORFs that can be found for each of our model organisms, and more than 270 for *E. coli* and *P. fluorescens*.

**Table 4:** Numbers of additional ORFs predicted by Prodigal that can be added to Ensembl gene annotations. Additional ORFs are chosen if there are no fewer than 50 nucleotides overlap with an Ensembl gene.

| Model Organism | Ensembl Genes | Add'l Prodigal ORFs |
| --- | --- | --- |
| *B. subtilis* | 4,011 | 62 |
| *C. crescentus* | 3,737 | 64 |
| *E. coli* | 4,052 | 270 |
| *M. genitalium* | 476 | 70 |
| *P. fluorescens* | 5,178 | 293 |
| *S. aureus* | 2,478 | 74 |

## Discussion

### *Ab initio* tools usually perform better than model-based

We found that *ab initio* tools usually perform better than model-based tools. While no one tool performed the best or worst across all metrics, the *ab initio* tools Prodigal, GeneMarkS-2, MetaGeneAnnotator, MetaGeneMark and GeneMarkS were ranked first to fifth respectively, across our 12 metric ranking (Figure 3).

Strains of the same species can exhibit large intraspecies variation (Van Rossum *et al*., 2020). Additionally, genes resulting from horizontal transfer, which is more frequent within species (Van Rossum *et al*., 2020), are likely to contain features from the donor strains which the rigid model-based methods are unable to recognise. GeneMark, a model-based tool, published in 1993, even when both target genome and model were *E. coli*, was identified as one of the worst performing tools in this study, possibly driven by the well-known large open pangenome of this species (Lukjancenko *et al*., 2010). The same was observed for *S. aureus*. While model-based tools can perform well even when their model and target genomes are different, in the case of Augustus, which when applied to the *C. crescentus* genome using the *S. aureus* model, it was only able to detect 3.93% EGS genes, but unexpectedly detected 78.75% when using the *H. sapiens* model. Unsurprisingly, model-based predictors have therefore fallen out of development and use over the last decade and *ab-initio* based tools such as Prodigal, GeneMarkS-2 and GLIMMER3 have become ubiquitous.

### Codon usage has a large influence on accuracy

We found that codon usage has a large influence on accuracy due to its influence on start and stop codon choice, even in model organisms.

The recoding of a stop codon as an amino acid is rare and seems to be taxa specific (Dybvig and Voelker, 1996). While many of the tools offered the ability to change codon tables (often accounting for TGA specifically), the correct codon tables or codon preferences for each genome cannot be known in advance of annotation of a novel organism. Despite this, we would expect that they should be able to predict a significant proportion of genes, even in the absence of the knowledge of a different codon usage table. Some tools such as Prodigal will assess a genome using both the universal and *Mycoplasma* translation table, however remarkably this did not increase the accuracy of the tool when analysing *M. genitalium* genome (see Figure 2). Overall TGA was never predicted as tryptophan-coding in this genome by any tool (see Supplementary Results 2).

While ATG is used for 80% of start codons in the canonical annotations for most prokaryote genomes, some species and even some species-spanning gene families have been shown to use very different start codon profiles (Villegas and Kropinski, 2008). The use of different start codons in prokaryote genomes has often been correlated to the genome-wide GC content: at extreme low and high GC (*<* 30% and *>* 80%), ATG and GTG respectively are often more prominent. In our study the extreme example of this was *C. crescentus* which uses ATG as a start codon only 69% of the time. This is likely driven by its GC content of 67%. All of the tools performed poorly at predicting the correct start codon in this species (Figure 2). This has been reported in the literature, specifically in relation to the lack of translation initiation sequence motifs traditionally used by prediction tools to identify the frame and start locus of a gene (Schrader *et al*., 2014). This is not unique to *C. crescentus* and as shown in Table 2, for all 6 model organisms incorrect start codon selection resulted in either elongated or truncated coding sequences (see Supplementary Results 2). The analysis of *E. coli* exhibited the lowest divergence between EGS and predicted start codon selection (see Supplementary Results 2 for more detail), possibly as a result of its historic use as a model organism and having the largest use of the canonical ATG start codon in this study. Studies continue to investigate the possible fluidity of gene start codon selection and how some genes recorded in genomic databases may either have been annotated with the wrong start codon, or even require the annotation of multiple alternative start positions and therefore start codons (Villegas and Kropinski, 2008; Meydan *et al*., 2019; Baranov *et al*., 2015).

### Metagenomic annotation approaches are suitable for whole genome sequences

Interestingly, tools made specifically for metagenomic and fragmented genome annotation performed better than most single genome tools (tools ranked 3rd, 4th and 6th were developed for metagenome annotation), possibly indicating that even ‘complete’ genomes may themselves still harbour elements of sequencing and assembly error which these types of algorithms have been designed to account for. Most genomes submitted to databases such as the NCBI Genome repository (Haft *et al*., 2017) are incomplete and can contain hundreds of fragments which can make gene prediction an even more difficult task. As S. Salzberg said in 2019 “Paradoxically, the incredibly rapid improvements in genome sequencing technology have made genome annotation less, not more, accurate.” (Salzberg, 2019). This indicates that future annotation work performed on non-model and more diverse organisms may benefit from approaches implemented by metagenomic tools.

### Short genes and overlapping genes are often misreported

We found that short ORFs and overlapping ORFs are often misreported and that many tools still have hard-coded limitations and weightings against these types of genes, with model-based tools performing especially poorly.

It has been well-established in the literature that Short genes are likely under-represented across genomic databases, and therefore, possibly even within the gold standard Ensembl data used in this study (Storz *et al*., 2014; Duval and Cossart, 2017; Su *et al*., 2013). The growing acceptance that Short genes are not only common in prokaryotic genomes but also have important roles (Andrews and Rothnagel, 2014), is at odds with many tools still containing hard-coded limitations for minimum ORF length and algorithmic weights against short ORFs. As might be expected because of its re-coding of TAG, *M. genitalium* proved challenging for all tools to accurately identify PCGs, resulting in the early truncation of a large proportion of genes and an increase in predicted short ORFs. This often led to the tools predicting additional spurious short ORFs in the missed regions. However, for the other genomes, most tools also predicted too many short ORFs (9.07% and 39.10%, for *ab initio* and model-based tools respectively), but paradoxically still managed to miss a large proportion of Short genes in the Ensembl annotations (missing 26.38% and 53.69% for *ab initio* and model-based tools respectively) (see Supplementary Table 6-8 and Results File 2).

For overlapping genes, while *ab initio* tools performed better than model-based tools (see Supplementary Tables 4-5), in general they both under-predicted the number of overlapping genes across the genomes (on average −6.07% and −30.15% for *ab initio* and model-based tools respectively) (see Supplementary Tables 4-5 and Results File 2). No tool was able to correctly detect more than 20% of *M. genitalium’s* overlapping EGS genes. Overlapping and nested genes have now become an area of renewed interest for their potential impact on genomic organisation and evolution (Huvet and Stumpf, 2014; Krakauer, 2000). For example, mokC in *E. coli*, believed to be a regulatory peptide, completely overlaps hokC and enables hokC expression (Pedersen and Gerdes, 1999) and no tool was able to detect both genes correctly.

Overall, the tools struggled to handle overlapping gene loci, and often returned either only one or neither of the overlapping coding regions in their predictions. This may be due to the manner in which many tools filter multiple candidate ORFs for a single locus leading to sub-optimal predictions. For example, Prodigal reports an ORF in *C. crescentus* on the positive strand at 23,760-24,074 when in fact the EGS-gene is 23,550-24,170 on the negative strand. The unallocated space (24,074-24,170) resulted in Prodigal reporting the next downstream ORF starting at 24,091, instead of 24,133 (as in the Ensembl annotation), erroneously including 5’ UTR in the predicted CDS.

There are now tools to identify putative short ORFs in both prokaryotes and eukaryotes (Bartholomaus *et al*., 2020; Ji *et al*., 2020). However, our results suggest that the identification of short ORFs and overlapping ORFs can not be done independently without consequences for annotation accuracy.

### Historic bias affects gene prediction today

Overall we have observed an increase in accuracy in tools over time as can be seen with the different versions of GeneMark compared here: the overall rankings of model-based GeneMark (1993) (with *E. coli/S. aureus* models), *ab initio* GeneMarkS (2001) and *ab initio* GeneMarkS-2 (2018) are 20/17, 5 and 2 respectively. However, GeneMarkS (2001) performed better than its successor GeneMarkS-2 (2018) for 5 out of the 12 metrics in Figure 3 including Percentage of Genes Detected (M1) in *P. fluorescens, M. genitalium* and *B. subtilis* (see Supplementary Results File 1). The performance of GeneMarkS (2001) in M1 may reflect its use for an extended period of time in the NCBI Prokaryote Annotation Pipeline. Possibly as a result of this, many of the genes GeneMarkS (2001) detected, were originally identified by the tool itself. Similarly, all model-based tools performed at their best across the 12 metrics and 6 model organisms when using their *E. coli* model, hinting at the impact of historical research in this organism. Advances in the realms of machine learning and statistical modelling have the greatest potential to address these issues but are also likely to be the most prone to historical biases in the databases. Many of the rules, such as standard ORF length and codon usage, are inferred from previously predicted PCGs. The existence of annotation errors and omissions in various sequence databases is well established and unlikely to be resolved in the near future (Bork and Bairoch, 1996; Karp, 1998) without significant coordination between repositories (Klimke *et al*., 2011). Additionally much of the sequence information is derived from model organisms will become of less relevance as greater numbers of novel organisms are sequenced (Hunter, 2008).

These issues have been raised previously: In 2009, the “Best Practices in Genome Annotation” meeting report listed a number of areas of concern put forward by attendees (Madupu *et al*., 2010) including tool choice, strategy to update and correct previous annotations, tracking of changes in databases, prioritisation of certain genes for experimental evaluation, documented processes and keeping up with technological advances. The work presented here addresses the issue of tool choice, but there is still much of the recommendations to be realised. The lack of any previous detailed systematic overview of method performance may also have played a part in these biases not being addressed to date.

## Conclusion

We have presented a comprehensive set of metrics which distinguish ORF prediction tools from each other and make it possible to identify which performs better for specific use-cases. The ORForise evaluation framework enables users to evaluate new and existing annotations and generate consensus and aggregate gene predictions. We have demonstrated that certain types of genes, such as short genes, overlapping genes and those with alternative codon usage, are still elusive, even to the most advanced *ab initio* techniques. Worryingly, the performance of any tool seems to depend on the genome that is being analysed. For instance, Prodigal which ranked best overall, was ranked first for *E. coli, S. aureus* and *P. fluorescens*, MetaGeneAnnotater was ranked first for *B. subtilis* and *M. genitalium* and GeneMarkS-2 was ranked first for *C. crescentus* (see Supplementary file Results 3). Additionally, no individual tool ranked as the most accurate across all genomes for the Percentage of Genes Detected (M1) (the single metric historically used to assess tool performance) or any other individual metric. This is likely to have a measurable impact on downstream genomic and pangenomic studies. However, overall we found Prodigal to be one of the most well-rounded tools, not only detecting the highest number of EGS-genes for two very diverse model organisms (*E. coli* and *M. genitalium*), but also performing overall best when ranked across the 12 metric rankings and 6 model organisms. It was also overall best for Perfect Matched genes (M5). However as outlined earlier, it was not always ranked first for all genomes, further suggesting that users should choose tools carefully, based on the organism and question they are studying. Finally, we advise against generating aggregated *ab initio* annotations from multiple tools where no gold standard exists, as this results in poor overall performance. However, additional cycles of annotation with tools designed to identify putative ORFs in the intergenic regions of gold standard annotations, show promise for improving current prokaryotic genomic knowledge.

## Supporting information

Supplemental Results 1 (Top 12 Metric Results)

Supplemental Results 2 (Full Metric Results)

Supplemental Rankings

## Author Contributions

All authors discussed the conceptualisation of the comparison platform and its impact. N.J.D wrote the code. All authors contributed to the manuscript.

## Footnotes

1 Available at https://github.com/NickJD/ORForise/tree/master/Genomes

