## Supplemental Document for "No one tool to rule them all: Prokaryotic gene prediction tool performance is highly dependent on the organism of study"

#### 1 Model Organisms (Ensembl Bacteria Release 46)

- *Bacillus subtilis (B. subtilis) - Strain BEST7003 - Assembly ASM52304v1: B. subtilis* is a Gram-positive, genetically tractable, non-pathogenic model organism used in the industrial production of enzymes. It is part of the Firmicute phylum and is a useful model in the study of *Mycobacterium tuberculosis*, which is the the causative agent of tuberculosis. The strain BEST7003 with assembly ASM52304v1 was chosen for this study (Itaya *et al.*, 2005).

- *Caulobacter crescentus (C. crescentus) - Strain CB15 - Assembly ASM690v1: C. crescentus* is a Gram-negative, oligotrophic bacterium commonly found throughout freshwater lakes and streams. It is an important model organism for studying the regulation of the cell cycle, asymmetric cell division, and cellular differentiation and is part of the Proteobacteria phylum. The CB15 strain with the ASM690v1 assembly was chosen for this study (Nierman *et al.*, 2001).

- *Escherichia coli (E. coli) K-12 - Strain ER3413 - Assembly ASM80076v1: E. coli* is one of the most extensively studied microorganisms and is part of the Proteobacterium phylum. *E. coli* is Gram-negative and its genome was first completely sequenced in 1997. It was chosen then for its unique biochemical, molecular and biotechnological attributes but it widely studied now due to its tractability. The K-12 ER3413 strain with the ASM80076v1 assembly was chosen for this study (Anton *et al.*, 2015).

- *Mycoplasma genitalium (M. genitalium) - Strain G37 - Assembly ASM2732v1: M. genitalium* is a parasitic bacterium with one of the smallest currently known genomes of any free living bacterium at around 580,000 bps. Due to it being a human pathogen and its unique genome size, *M. genitalium* has been used as a model for a minimal organism in the study of essential genes due to being one of the most streamlined bacterial genomes currently known (Glass *et al.*, 2006). Although *M. genitalium* does not have cell walls, it is believed to have evolved from Gram-positive bacteria which had lost their cell wall and is part of the Firmicute phylum. The G-37 strain with ASM2732v1 assembly was chosen for this study (Hutchison *et al.*, 1999).

- *Pseudomonas fluorescens (P. fluorescens) - Strain UK4 - Assembly ASM73042v1: P. fluorescens* is a rod-shaped, Gram-negative bacterium and is part of the Proteobacteria phylum.The antibiotic Mupirocin can be produced by cultured *P. flurescens* and is used in the treatment of skin, ear and eye disorders and is a model organism for cell cycle, cell division and differentiation. The UK4 strain with the ASM73042v1 assembly was chosen for this study (Dueholm *et al.*, 2014).

- *Staphylococcus aureus ( S. aureus) - Strain 502A - Assembly ASM59796v1: S. aureus* is Gram-positive bacterium of the Firmicute phylum and is commonly found on the human body, including the nose, skin and the respiratory tract. It has been known to cause diseases such as infective endocarditis and a drug resistant strain is commonly known as Methicillin-resistant *Staphylococcus aureus* (MRSA). The 502A strain with assembly ASM59796v1 was chosen for this study (Parker *et al.*, 2014).

### 2 Supplementary Figures

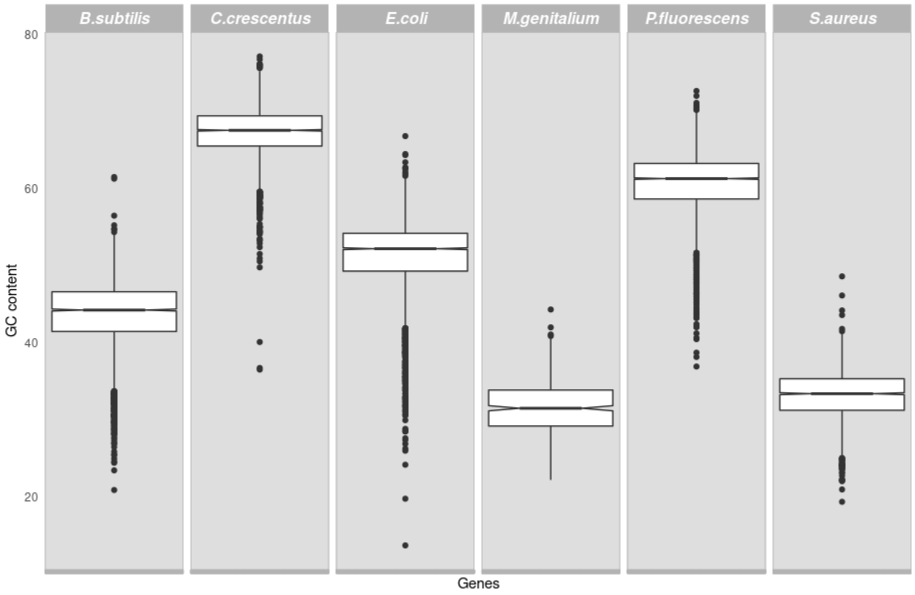


Figure 1: GC content of the six model organisms and their Ensembl annotated protein coding genes. Note the high levels of variance within and between each genome.

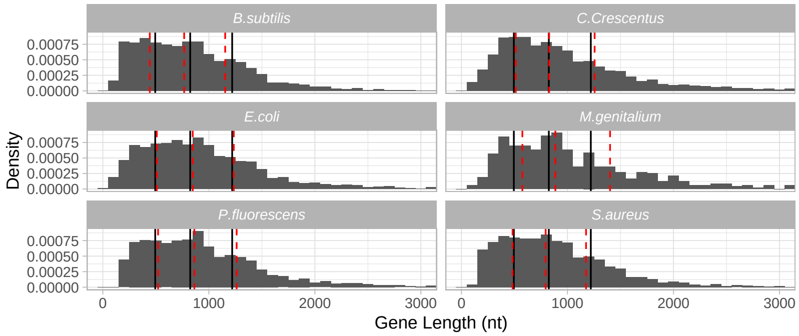


Figure 2: Protein-Coding Gene (PCG) lengths plotted for each model organism. The black, solid vertical lines are at the overall first quartile (494), median (824) and third quartile (1220) for all six model model organisms. The red dotted lines show the first quartile, the median and the third quartile for each organism individually. The x-axis is truncated at 3000 nt. The proportion of PCG lengths at or below this value are 0.964 for *M. genitalium*, 0.984 for *P. fluorescens*, 0.987 for *E. coli*, *S. aureus* and *C. crescentus*, and 0.990 for *B. subtilis*. A total of 23 PCGs were longer than 5000 nt. The distributions of PCG lengths for *E. coli*, *S. aureus*, *C. crescentus* and *P. fluorescence* are comparable to the overall distribution. The lengths for *B. subtilis* are somewhat smaller than expected overall, while the lengths for *M. genitalium* are longer than expected.

### 3 Prediction Tools

#### 3.1 Prediction Tools Run-Parameters

All tools were provided with the same 6 DNA data files each containing the complete genome for an organism in a single sequence. The model-based tools were provided with two different organism models, *E. coli* K-12 and *S. aureus* - Mu50 (strains selected where possible). These were chosen as both were in the set of six bacteria and were models which were already available for all model-based tools. In addition, Augustus, which was originally developed for eukaryotic gene prediction, was run with the inclusion of the *H. sapiens* model and each individual Coding Sequence (CDS) predicted was taken as a PCG. We did not provide genome-specific parameters such as alternative codon tables to the tools as this study aimed to be representative of real-world analysis where such information may not be known. Each tool was run using its default parameters with no user-defined filtering.

This was performed locally on a 64-bit Linux machine with an i7 2600k CPU with 32GB RAM and none of the tools required more than a few minutes to run or more than 500 MB of RAM.

While most of the tools were available as online resources, they were downloaded from the links included in their associated publications and the specific versions used are listed below.

#### 3.2 Prediction Tools

- Model-based group:

  Some tools have been designed for a specific set genomes or strains and require a pre-built model (a rigid set of parameters tuned to a particular organism) to perform predictions. The construction of these models rely heavily on having an accurate and complete set of genes for a particular organism (among other information). While inaccuracies or biases in the data are likely to be present in the final models, model-based gene predictors trained on a particular species are expected to perform well on strains with comparable gene and genome structure. Overfitting can occur, where only similar genes to those previously found are detected at a high sensitivity. However, there can be large differences in gene number, gene length and genome size between strains of the same species. Model-based prediction for certain model organisms where specific strains are often used for scientific and industrial purposes can still be effective as there may be little genetic difference between two isolates of the same strain.

  *E. coli* **K12** and *S. aureus* **Mu50** (strain-specific where possible) were chosen as prediction models for each of the prediction tools below. Prediction with Augustus was also performed with the Eukaryote *H. sapiens* model, and when the model predicted multiple introns for a gene we took each predicted CDS as an ORF.

  - **Augustus** Keller *et al.* (2011) - Version 3.3.3
    Originally published in 2003, Augustus was developed as a eukaryote genome prediction tool combining protein-family-based gene prediction and incorporated knowledge from external sources (pre-computed genome models) to combine them with an *ab initio* prediction to specifically help with exon prediction. Later versions of Augustus included 3 bacterial and 1 archaeal species to the pre-computed model list to allow for a selection of prokaryotic genome annotation.
  - **EasyGene** Nielsen and Krogh (2005) - Version 1.2
    EasyGene 1.2 published in 2005, employs a genome specific Hidden Markov Model (HMM) which after extracting all ORFs above 120 nt, filters them by using a sequence similarity search to a protein database. The resulting genes and their start positions are then used to retrain the HMM. EasyGene produces scores for multiple potential start codons for each gene and selects the one with the highest computed confidence value.
  - **GeneMark.hmm** Lukashin and Borodovsky (1998) - Prokaryote Model Version 3.2.5
    GeneMark.hmm, published in 1998 was developed to be one of the first tools to "improve the gene prediction quality in terms of finding exact gene boundaries". A HMM is used to model gene boundaries as transitions between hidden states along with ribosomal binding site patterns to refine translation initiation codons. The current genome model parameters were derived from the use of GeneMarkS, the successor of GeneMark.hmm.
  - **GeneMark** Borodovsky and McIninch (1993) - Version 2.5
    GeneMark, developed in 1993, was one of the first gene prediction methods to efficiently perform

whole-genome annotation, notably for its ability to predict ORFs on both strands of DNA simultaneously. Markedly, GeneMark was used for the first annotation of a completely sequenced bacterium, *Haemophilus influenzae*, and the first completely sequenced archaeon, *Methanococcus jannaschii.* The GeneMark algorithm consists of species-specific inhomogeneous Markov chain models computed from protein-coding DNA sequences and homogeneous Markov chain models of non-coding DNA. Probability of a predicted sequence fragment to be protein coding in one of six possible frames (including three frames in complementary DNA strand) or to be "non-coding" is computed to determine potential genes in the opposite strand of DNA.

  - **FGENESB** Salamov and Solovyevand (2011) - The FGENESB pipeline identifies protein, tRNA and rRNA genes, potential promoters, terminators and operons and performs an initial prediction of 'long' ORFs as a starting point for calculating parameters for gene prediction. The gene prediction algorithm is based on Markov chain models of coding regions and their translation and termination sites. Furthermore, operon prediction is performed using distances between ORFs, frequencies of neighboring genes in known bacterial genomes and positions of predicted promoters and terminators. FGENESB, unlike other model-based prediction tools, presents its model selection as "Choose closest organism", rather than "select species/organism", indicating that the developers acknowledge the models may be used as best-fit rather than for exact species prediction.

- *Ab initio* group:

  Self-training tools do not require any previous knowledge of the target genome and predict *ab initio*, directly from sequence. Tools such as Prodigal were developed to be used on different prokaryotic organisms, however, they do rely on broad models either trained on features gathered directly from the input genome or predict ORFs using a set of predefined parameters which may be adapted. The criteria considered while making predictions include but are not limited to, overlapping ORFs, GC content, ORF length, predicted start and stop codons, and distances between ORFs Delcher *et al.* (1999); Besemer *et al.* (2001). Unfortunately, these criteria and their thresholds are still based on prior knowledge as deciding between candidate ORFs still requires a number of assumptions based on previously studied genes and genomes which the developer has embedded into the algorithm.

  - **Prodigal** Hyatt *et al.* (2010) - Version 2.6.2 Prodigal is an unsupervised gene predictor which examines the input genome for the creation of its input-specific training set. 100 prokaryote genomes were selected in the initial development of the algorithm to determine "very general rules about the nature of prokaryotic genes, such as gene size, maximum overlap between two genes... and RBS (ribosomal binding site) motif usage". A number of constants within the algorithm were tuned to the genetic makeup of the 100 genomes. GC is an important statistic for Prodigal and it is used for a number of steps in the prediction process such as coding scores for each gene predicted. Prodigal performs a number of scoring functions on different aspects of each DNA region selected, thus producing a set of putative "most-likely real" genes. These genes are then examined and are used to tune the model before prediction of genes which exhibit lower likelihood scores. Furthermore, Prodigal has been designed to detect whether genetic code 4 is needed (*Mysoplasma*) and use it instead of the default code 11.
  - **GeneMarkS** Besemer *et al.* (2001) - Version 4.25 Developed in 2001, GeneMarkS was one of the first *ab initio* gene prediction methods which could learn directly from short (>400) sequences without prior knowledge or pre-trained models. As with other contemporary tools, HMMs were trained on protein-coding sequence data, non-coding DNA samples and modelled on transition and initiation parameters trained from input sequence. Codon frequencies and positional statistics are utilised along with genomic GC content to learn coding potential for identified ORFs. GeneMarkS has become a bedrock for future prediction tools and has been used as part of wider genome annotation pipelines.
  - **GeneMarkS 2** Lomsadze *et al.* (2018) - Version '2020' An advancement over the original GeneMarkS tool, GeneMarkS-2 further utilises a self-derived *ab initio* training model learnt from input sequences for finding species-specific (native) genes. A collection of pre-computed "heuristic" models are utilised to identify harder-to-detect genes (horizontally transferred). GeneMarkS-2 learns distinct sequence patterns inherent to prokaryotic genomes which are involved in gene expression control. The majority of protein-coding regions in prokaryotic genomes are known to carry species-specific codon usage patterns and GeneMarkS-2 learns these patterns and estimates parameters of typical protein-coding

regions of a target genome. This process is similar to the one employed by GeneMarkS(1) but extended.

- **GLIMMER 3** Delcher *et al.* (2007) - Version 3.02 GLIMMER 3, published in 2007 is the third iteration of the GLIMMER microbial gene predictor software. A number of improvements over the previous implementations include improved coding region and start codon detection, along with a reduction in incorrectly reported overlapping genes. GLIMMER 3, as with the previous versions, starts by predicting ORFs with little filtering and then using a number of user defined (or default) parameters (such as start codon selection, ORF and overlap length). These ORFs are then scored for their coding potential. To overcome the high levels of potential false positive overlapping ORFs, GLIMMER3 uses these scores to select which of any two overlapping ORFs are more likely to be real (in cases where maximum overlap is surpassed). An Interpolated Markov Model (IMM) is used in the prediction process to help identify coding regions and has also been shown to separate DNA between bacterium and host DNA. GLIMMER, along with GeneMarkS, was also used as part of the NCBI prokaryote annotation pipeline (Tatusova *et al.*, 2016).
- **GeneMark Heuristic Approach** Besemer and Borodovsky (1999) - Version 3.25 As with many other tools from the GeneMark suite, GeneMark Heuristic Approach (GeneMark HA) was developed on the observations made from GeneMark and GeneMark.hmm. The method was designed to build Markov models derived on a minimal amount of DNA information from 17 completed bacterial genomes. Linear regression was performed to approximate relationships between positional and global nucleotide frequencies, relationships between the amino acid frequencies and the global GC% of the bacterial genomes. Amino acids frequencies were calculated mostly from an *E. coli* genome to build constants for the algorithm. The algorithm builds a heuristic model for every sequence longer than 400 nt. GeneMark HA derived models are expected to be applied to the analysis of the input sequence by the GeneMark and GeneMark.hmm programs.
- **TransDecoder** Haas *et al.* (2013) - TransDecoder was designed to identify candidate coding regions within transcript sequences, such as those generated by *de novo* RNA-Seq transcript assembly, or constructed based on RNA-Seq alignments to the genome. TransDecoder identifies likely coding sequences based on a minimum ORF length and a computed log-likelihood score >0. The coding score is greatest when the ORF is scored in the 1st reading frame as compared to scores in the other 5 reading frames. The longer of two ORFs is reported if one is encapsulated by the others coordinates. However, a single transcript can report multiple ORFs (allowing for operons, chimeras, etc).
- **FragGeneScan** Rho *et al.* (2010) - FragGeneScan has been specifically designed to improve prediction performance on metagenomic and short-read sequence data with high levels of sequencing errors, but also perform comparably with other contemporary tools on complete genomes. A combination of probabilistic models trained on codon usage and sequence error data, was used to evaluate sections of DNA for their gene encoding potential. This method has shown higher performance for predicting genes on short-reads with high levels of sequence error than other contemporary methods but can be used on complete low-error genomes.
- **MetaGene** Noguchi *et al.* (2006) - MetaGene, one of the first ORF prediction tools specifically developed for prediction on fragmented and metagenomic genomes, examines di-codon frequencies estimated by the GC content of a given sequence with other measures such as length, distance to between ORFs and start codon distribution. MetaGene can predict a whole range of prokaryotic genes based on the anonymous genomic sequences of a few hundred bases and identify partial ORFs which have are located on the terminus of the fragmentary genomic sequences.
- **MetaGeneMark** Zhu *et al.* (2010) - The hueristic model behind MetaGeneMark was developed to replace traditional methods of ORF prediction parameter estimation such as supervised training on a set of "validated" genes or unsupervised training on an input sequence. Dependencies which had formed in evolution, between codon frequencies and genome nucleotide composition are utilised to derive patterns of codon frequencies, critical for the model parameterisation, from frequencies of nucleotides observed in a short or metagenomic sequences. An effective method to estimate prediction parameters was derived from the frequencies of oligonucleotides in protein-coding regions and whole-genome nucleotide composition.
- **Meta Gene Annotator** Noguchi *et al.* (2008) - Published in 2008, MetaGeneAnnotator predicts all kinds of prokaryotic genes from anonymous genomic sequences. It integrates statistical models

of prophage, bacterial and archaeal genes, and builds a self-trained model from input sequences for the predictions. This results in the detection of not only "typical genes but also atypical genes, such as horizontally transferred and prophage genes in a prokaryotic genome". The algorithm also includes a novel approach for the analysis of ribosomal binding sites, which has enabled the detection of species-specific patterns, thus allowing for "precise" prediction of translation starts sites.

### 4 Code and analysis scripts

To inspect the various differences between the six genomes, a number of additional Python3 scripts were written to interrogate the canonical annotations. These scripts are available at `https://github.com/NickJD/ORForise/tree/master/Post_Analysis_Scripts`).

Python3 (Van Rossum and Drake, 2009) with Matplotlib (Hunter, 2007) and R (R Core Team, 2020) with ggplot2 (Wickham, 2016) were used to produce the figures.

### 5 Description of Comparison Metrics

- Number of ORFs:
  This is the number of ORFs which the tool has predicted. Some tools predict a large number of potential ORFs and then filter them. This metric corresponds to the remaining ORFs presented to the user after default filtering.
- Percentage Difference of All ORFs: **(M3)**
  This is the percentage change between the number of predicted ORFs and the number of actual Ensembl Gold Standard Genes. *100 * (Number of ORFs - Number of Genes) / Number of Genes*
- Number of ORFs that Detect a Gene:
  This is the number of ORFs which correctly detect at least 75% of the nucleotides of an Ensembl Gold Standard Gene and are in the same frame.
- Percentage of ORFs that Detected a Gene: **(M2)**
  This is the percentage of ORFs which correctly detect at least 75% of the nucleotides of an Ensembl Gold Standard Gene and are in the same frame.
- Number of Genes Detected:
  The number of Ensembl Gold Standard Gene Detected is reported separately here as one ORF can cover or overlap with one or more smaller genes. The minimum 75% of nucleotide detection and correct frame still apply.
- Percentage of Genes Detected: **(M1)**
  The percentage of Ensembl Gold Standard Gene Detected is reported separately here as one ORF can cover or overlap with one or more smaller genes. The minimum 75% of nucleotide detection and correct frame still apply.
- Median Length of All ORFs:
  Median length of all predicted ORFs in nucleotides.
- Median Length Difference: **(M4)**
  This is the Percentage Difference from the mean length of Ensembl Gold Standard Genes compared to the mean length of all predicted ORFs. *100 * (Median ORF Length - Gene Median Length) / Gene Median Length*
- Minimum Length of All ORFs:
  The length of the shortest predicted ORF in nucleotides.
- Minimum Length Difference:
  This is the percentage difference from the shortest Ensembl Gold Standard Gene compared to the length of the shortest predicted ORF. *100 * (Minimum ORF Length - Minimum Gene Length) / Minimum Gene Length*
- Maximum Length of All ORFs:
  The length of the longest predicted ORF in nucleotides.
- Maximum Length Difference:
  This is the percentage difference from the longest Ensembl Gold Standard Gene compared to the length of the longest predicted ORF. *100 * (Maximum ORF Length - Maximum Gene Length) / Maximum Gene Length*
- Median GC Content of all ORFs:
  This median GC content calculated from all predicted ORFs.
- Percentage Difference of all ORFs Median GC:
  This is the Percentage Difference of the median GC content of all predicted ORFs compared to the median GC content of all Ensembl Gold Standard Genes. *100 * (Median GC content of all ORFs - Median GC content of all Genes) / Median GC content of all Genes*

- Median GC Content of Matched ORFs:
  This median GC content calculated from predicted ORFs which detected an Ensembl Gold Standard Gene.

- Percentage Difference of Matched ORF GC:
  This is the Percentage Difference of the median GC content of ORFs which detected an Ensembl Gold Standard Gene compared to the median GC content of all Ensembl Gold Standard Genes. *100 * (Median GC content of Matched ORFs - Median GC content of all Genes) / Median GC content of all Genes*

- Number of ORFs which Overlap Another ORF:
  This is the number of predicted ORFs which overlap another predicted ORF by at least one nucleotide base.

- Percentage Difference of Overlapping ORFs:
  This is the Percentage Difference of overlapping ORFs as compared to the number of overlapping Ensembl Gold Standard Genes. *100 * (Number of Overlapping ORFs - Number of Overlapping Genes) / Number of Overlapping Genes*

- Maximum ORF Overlap:
  This is the maximum length of ORF overlap in nucleotides.

- Median ORF Overlap:
  This is the median length of ORF overlap calculated from all ORF overlap lengths.

- Number of Matched ORFs Overlapping Another ORF:
  This is the number of ORFs which detected a gene that overlap another predicted ORF by at least one base.

- Percentage Difference of Matched ORFs Overlapping another ORFs: **(M8)**
  This is the percentage difference of overlapping ORFs which detected a gene as compared to the number of overlapping annotated genes. *100 * (Number of Matched Overlapping ORFs - Number of Overlapping Genes) / Number of Overlapping Genes*

- Maximum MatchedORF Overlap:
  This is the maximum length of matched ORF overlap in nucleotides.

- Median Matched ORF Overlap:
  This is the median length of matched ORF overlap calculated from all ORF overlap lengths.

- Number of Short-ORFs:
  This is the number of predicted ORFs which are under 100 nucleotide bases.

- Percentage Difference of Short-ORFs:
  This is the percentage difference of predicted Short-ORFs as compared to the number of annotated Short-Genes. *100 * (Number of Short-ORFs - Number of Short Genes) / Number of Short Genes*

- Number of Short-Matched-ORFs: **(M9)**
  This is the number of ORFs which detected an annotated gene and which are under 100 nucleotide bases.

- Percentage Difference of Short-Matched-ORFs:
  This is the percentage difference of Short-ORFs which detected a gene as compared to the number of annotated Short-Genes. *100 * (Number of Short-Matched-ORFs - Number of Short Genes) / Number of Short Genes*

- Number of Perfect Matches: **(M5)**
  This is the number of ORFs which have correctly identified the exact start and stop position of an annotated gene.

- Percentage of Perfect Matches:
  This is the percentage of ORFs which have correctly found an annotated gene and have identified both the exact start and stop position. *100 * Number of ORFs which Matched a Gene - Number of Genes) / Number of Genes*

- Number of Perfect Starts:
  This is the number of Matched ORFs which have correctly identified the start position of an annotated gene.

- Percentage of Perfect Starts:
  This is the percentage of Matched ORFs which have correctly identified an annotated gene and its start position.

- Number of Perfect Stops:
  This is the number of Matched ORFs which have correctly identified the stop position of an annotated gene.

- Percentage of Perfect Stops:
  This is the percentage of Matched ORFs which have correctly identified an annotated gene and its stop position.

- Number of Out of Frame ORFs:
  This is the number of ORFs which locus covered more than 75% of an annotated gene but were out of frame, thus classified as Unmatched.

- Number of Matched ORFs Extended a Coding Region:
  This is the number of Matched ORFs which locus extend the 3 and 5-prime end of its identified annotated gene.

- Percentage of Matched ORFs Extended a Coding Region:
  This is the percentage of Matched ORFs which locus extend the 3 and 5-prime end of its identified annotated gene.

- Number of Matched ORFs Extended Start Region:
  This is the number of Matched ORFs which locus extend the 5-prime end of its identified annotated gene.

- Percentage of Matched ORFs Extended a Coding Region:
  This is the percentage of Matched ORFs which locus extend the 5-prime end of its identified annotated gene.

- Number of Matched ORFs Extended Stop Region:
  This is the number of Matched ORFs which locus extend the 3-prime end of its identified annotated gene.

- Percentage of Matched ORFs Extended a Coding Region:
  This is the percentage of Matched ORFs which locus extend the 3-prime end of its identified annotated gene.

- Number of All ORFs on Positive Strand:
  This is the number of all predicted ORFs on the positive strand.

- Percentage of All ORFs in Positive Strand:
  This is the percentage of all predicted ORFs on the positive strand.

- Number of All ORFs in Negative Strand:
  This is the number of all predicted ORFs on the negative strand.

- Percentage of All ORFs in Negative Strand:
  This is the percentage of all predicted ORFs on the negative strand.

- Mean Start Difference of Matched ORFs: **(M6)**:
  This is the mean difference calculated by taking all matched ORF start position prediction differences from the identified annotated genes and applying a median calculation. This is calculated in nucleotides and the closer to 0, the lower the difference or effective error.

- Mean Stop Difference of Matched ORFs: **(M7)**
  This is the mean difference calculated by taking all matched ORF stop position prediction differences from the identified annotated genes and applying a median calculation. This is calculated in nucleotides and the closer to 0, the lower the difference or effective error.
- ATG Start Percentage:
  This is the percentage of all predicted ORFs which begin with the ATG codon. .
- GTG Start Percentage:
  This is the percentage of all predicted ORFs which begin with the GTG codon.
- TTG Start Percentage:
  This is the percentage of all predicted ORFs which begin with the TTG codon.
- ATT Start Percentage:
  This is the percentage of all predicted ORFs which begin with the ATT codon.
- CTG Start Percentage:
  This is the percentage of all predicted ORFs which begin with the CTG codon.
- Other Start Codon Percentage:
  This is the percentage of all predicted ORFs which begin with an alternative start codon.
- TAG Stop Percentage:
  This is the percentage of all predicted ORFs which begin with the TAG codon.
- TAA Stop Percentage:
  This is the percentage of all predicted ORFs which begin with the TAA codon.
- TGA Stop Percentage:
  This is the percentage of all predicted ORFs which begin with the TGA codon.
- Other Stop Codon Percentage:
  This is the percentage of all predicted ORFs which end with an alternative stop codon.
- True Positive:
  The true positive value is calculated by dividing the number of annotated genes correctly identified by the total number of annotated genes(75% identified and in frame). *Number of Genes Detected / Number of Genes*
- False Positive:
  The false positive value is calculated by dividing the number of predicted ORFs which did not match any annotated genes by the total number of annotated genes.*Number of Unmatched ORFs / Number of Genes*
- False Negative:
  The false negative value is calculated by dividing the number of Ensembl genes missed by the predicted ORFs by the total number of Ensembl genes.
- Precision: **(M10)**
  The precision value is calculated by dividing the true positive value by the true positive and false positive values combined.
- Recall: **(M11)**
  The recall value is calculated by dividing the true positive value by the true positive and false negative values together.
- False Discovery Rate: **(M12)**
  The false discovery rate is calculated by dividing the false positive value by the false positive and true positive values combined.

- True Positive (Nucleotide):
  The true positive value is calculated by dividing the number of nucleotides in Ensembl genes correctly identified by the total number of nucleotides in all Ensembl genes.
- False Positive (Nucleotide):
  The false positive value is calculated by dividing the number of nucleotides in predicted ORFs which were not in any Ensembl genes by the total number of nucleotides in all Ensembl genes.
- True Negative (Nucleotide):
  The true negative value is calculated by dividing the number of nucleotides not in any ORFs by the number of nucleotides
- False Negative (Nucleotide):
  The false negative value is calculated by dividing the number of nucleotides in Ensembl genes incorrectly identified by the total number of nucleotides in all Ensembl genes.
- Precision (Nucleotide):
  This precision value is calculated by dividing the nucleotide true positive value by the nucleotide true positive and false positive values combined.
- Recall (Nucleotide):
  This recall value is calculated by dividing the nucleotide true positive value by the nucleotide true positive and false negative values together.
- False Discovery Rate (Nucleotide):
  This false discovery rate is calculated by dividing the nucleotide false positive value by the nucleotide false positive and true positive values combined.
- ORF Nucleotide Coverage of Genome:
  This is the percentage of nucleotides in all predicted ORFs out of all nucleotides in the genome.
- Correctly Matched ORF Nucleotide Coverage of Genome:
  This is the percentage of nucleotides in Matched ORFs which correctly detected an Ensembl gene out of all nucleotides in the genome.

#### 6 Supplementary Tables

Table 1: Start codon usage for Ensembl gold standard protein coding genes for the six model organisms. Note the variation in usage of canonical start codon ATG and the alternative GTG and TTG codons.

| Model Organism | ATG | GTG | TTG | ATT | CTG | Other |
|---|---|---|---|---|---|---|
| *B. subtilis* | 76.81% | 10.10% | 13.09% | 0.00% | 0.00% | 0.00% |
| *C. crescentus* | 68.58% | 17.69% | 13.73% | 0.00% | 0.00% | 0.00% |
| *E. coli* | 90.67% | 7.50% | 1.70% | 0.05% | 0.05% | 0.02% |
| *M. genitalium* | 88.45% | 7.56% | 3.99% | 0.00% | 0.00% | 0.00% |
| *P. fluorescens* | 88.55% | 7.55% | 2.92% | 0.21% | 0.48% | 0.29% |
| *S. aureus* | 86.80% | 6.62% | 6.58% | 0.00% | 0.00% | 0.00% |

Table 2: Stop codon usage for Ensembl gold standard protein coding genes for the six model organisms. *M. genitalium* recodes TGA for Tryptophan and *E. coli* uses CTT for one gene.

| Model Organism | TAG | TAA | TGA | Other |
|---|---|---|---|---|
| *B. subtilis* | 13.96% | 62.93% | 23.11% | 0.00% |
| *C. crescentus* | 32.78% | 19.86% | 47.36% | 0.00% |
| *E. coli* | 6.89% | 64.68% | 28.41% | 0.02% |
| *M. genitalium* | 27.10% | 72.90% | 0.00% | 0.00% |
| *P. fluorescens* | 14.18% | 30.42% | 55.41% | 0.00% |
| *S. aureus* | 15.01% | 74.17% | 10.82% | 0.00% |

Table 3: GC content differences for Prodigal annotations. Shown here as median values are; GC content of Ensembl gold standard protein coding genes, the genes detected by Prodigal, those Prodigal obtained a partial match and those it missed.

| Model Organism | Ensembl GC | Detected GC | Partial GC | Missed GC |
|---|---|---|---|---|
| *B. subtilis* | 44.19% | 44.25% | 43.99% | 39.13% |
| *C. crescentus* | 67.52% | 67.71% | 67.69% | 65.65% |
| *E. coli* | 52.15% | 52.21% | 52.14% | 43.14% |
| *M. genitalium* | 31.44% | 32.90% | 32.75% | 30.76% |
| *P. fluorescens* | 61.25% | 61.36% | 60.25% | 53.36% |
| *S. aureus* | 33.33% | 33.33% | 30.13% | 32.62% |

Table 4: Percentages of the Ensembl gold standard protein coding genes and ORFs identified as overlapping. We show averages for *ab initio* and model-based predicted ORFs.

| Model Organism | Ensembl | *Ab initio* | Model-Based |
|---|---|---|---|
| *B. subtilis* | 21.37% | 21.44% | 15.44% |
| *C. crescentus* | 32.73% | 25.51% | 21.84% |
| *E. coli* | 22.53% | 22.68% | 18.20% |
| *M. genitalium* | 46.43% | 16.47% | 11.65% |
| *P. fluorescens* | 24.16% | 25.42% | 18.08% |
| *S. aureus* | 19.61% | 19.98% | 15.72% |

Table 5: Percentage Difference of overlapping ORFs as compared to the Ensembl gold standard protein coding genes. *Ab initio* and model based tools are separated into 2 groups each. 'Matched' represents the Percentage Difference for those ORFs which were able to detect an Ensembl gold standard protein coding gene whereas 'All' represents the Percentage Difference of the number of overlapping ORFs across all predicted ORFs.

| Group | Average | Standard Deviation | Standard Error |
|---|---|---|---|
| Matched, *ab initio* | -23.62% | 7.16% | 2.27% |
| Matched, model | -52.89% | 24.79% | 7.47% |
| All, *ab initio* | -6.07% | 11.55% | 3.65% |
| All, model | -30.15% | 29.41% | 8.87% |

Table 6: Percentage of the Ensembl gold standard protein coding genes and ORFs categorised as Short ORFs ($\geq$ 300 nt). We show averages for *ab initio* and model-based predicted ORFs. Note the large increase in Short ORFs predicted for *M. genitalium.*

| Model Organism | Ensembl | *Ab initio* | Model-based |
|---|---|---|---|
| *B. subtilis* | 13.66% | 12.58% | 13.24% |
| *C. crescentus* | 7.60% | 8.11% | 15.33% |
| *E. coli* | 10.24% | 10.45% | 13.04% |
| *M. genitalium* | 4.83% | 38.44% | 36.99% |
| *P. fluorescens* | 7.84% | 9.06% | 19.01% |
| *S. aureus* | 10.05% | 11.26% | 15.59% |

Table 7: Percentage Difference of short ORFs ($\leq$100 amino acids) as compared to the Ensembl gold standard protein coding genes. *Ab initio* and model based tools are separated into 2 groups each. 'Matched' represents the Percentage Difference for those ORFs which were able to detect an Ensembl gold standard protein coding gene whereas 'All' represents the Percentage Difference of the number of Short ORFs across all predicted ORFs. The results from *M. genitalium* were not included in this table's calculations.

| Group | Average | Standard Deviation | Standard Error |
|---|---|---|---|
| Matched, *ab initio* | -26.38% | 25.68% | 8.12% |
| Matched, model | -53.69% | 21.71% | 6.55% |
| All, *ab initio* | 9.07% | 39.87% | 12.61% |
| All, model | 39.10% | 91.22% | 27.50% |

Table 8: *M. genitalium*-only Percentage Difference of short ORFs ($\leq$100 amino acids) as compared to the Ensembl gold standard protein coding genes. *Ab initio* and model based tools are separated into 2 groups each. 'Matched' represents the Percentage Difference for those ORFs which were able to detect an Ensembl gold standard protein coding gene whereas 'All' represents the Percentage Difference of the number of Short ORFs across all predicted ORFs.

| Group | Average | Standard Deviation | Standard Error |
|---|---|---|---|
| Matched, *ab initio* | -27.34% | 25.15% | 7.95% |
| Matched, model | -55.28% | 20.62% | 6.22% |
| All, *ab initio* | 261.11% | 139.28% | 44.04% |
| All, model | 148.00% | 164.58% | 49.62% |

Table 9: Numbers of Ensembl genes which form an intersection (100% or 75%) with ORFs predicted by Prodigal.

| Model Organism | Ensembl Genes | Prodigal ORFs | 100% Intersection | 75% Intersection |
|---|---|---|---|---|
| *B. subtilis* | 4,011 | 4,016 | 3,673 | 3,943 |
| *C. crescentus* | 3,737 | 3,704 | 2,393 | 3,433 |
| *E. coli* | 4,052 | 4,263 | 3,737 | 3,973 |
| *M. genitalium* | 476 | 995 | 128 | 190 |
| *P. fluorescens* | 5,178 | 5,421 | 4,736 | 5,100 |
| *S. aureus* | 2,478 | 2,534 | 2,434 | 2,457 |
